# Putative biosurfactants from polar microbial consortia

**DOI:** 10.64898/2026.11.02.763558

**Authors:** Thamali Vidanage, Durell Desmond, R. Eric Collins

## Abstract

Biosurfactants (BS) are amphiphilic microbial metabolites that are biodegradable, low in toxicity, and environmentally compatible, making them attractive for applications in food, pharmaceutical, and bioremediation industries. This study investigated the phylogenetic composition and biosurfactant-producing potential of marine oil snow (MOS)-associated microbial consortia from the Churchill Marine Observatory, western Hudson Bay. MOS aggregates were cultivated in mineral Bushnell–Haas (BH) medium supplemented with 3% (w/v) NaCl, using motor oil as the sole hydrocarbon source. Enrichment cultures capable of growth in BH medium were subsequently evaluated for biosurfactant production using qualitative and quantitative assays at 4 °C and 20 °C. All cultures were maintained as enrichment consortia throughout the study.

A total of eight enrichment cultures were identified as biosurfactant-producing consortia. 16S rRNA gene sequencing revealed that *Pseudoalteromonas* dominated all consortia, while *Paraglaciecola, Marinomonas*, and other taxa were present at varying relative abundances. Although none of the consortia exhibited high emulsification activity (E_24_ > 40%), biosurfactant yields were significantly higher at 4 °C. Additionally, three consortia tested positive for rhamnolipid-type biosurfactant production.

## 1 Introduction

Biosurfactants (BS) can reduce fuel–water interfacial tension to form emulsions that facilitate the assimilation of hydrocarbons by microorganisms (Ruiz et al., 2021). Chemical and microbial surfactants are amphiphilic compounds, meaning that they contain both hydrophobic and hydrophilic groups. The hydrophilic groups consist of polysaccharides, peptides, or proteins, while the hydrophobic groups are made up of hydroxylated fatty acids or fatty alcohols (Lang, 2002). Bioemulsifiers (BE) are high-molecular-weight molecules such as lipoproteins, lipopolysaccharides, proteins, and exopolysaccharides (EPS) (Collins and Margesin, 2019). Both BS and BE are bioproducts produced by microorganisms that are less toxic than their chemical counterparts (Banat et al., 2014). The physical and chemical properties of biosurfactants make them suitable for various applications, including bioremediation of contaminated sites, microbial enhanced oil recovery, and use in the pharmaceutical, cosmetic, and food industries (de Lemos et al., 2023). Using biosurfactants as a potential oil remediation method has gained a lot of interest due to their effectiveness, environmental safety, and sustainability. Even though short-chain hydrocarbons such as alkanes and alkenes are readily biodegradable by many bacteria, long-chain petroleum hydrocarbons’ biodegradation takes a long time. Hence, the removal of those complex hydrocarbons from oil spills is challenging (Kubota et al., 2008).

The amphiphilic properties of biosurfactants provide a significant advantage by allowing them to spontaneously separate into two immiscible phases. This occurs as they reduce surface tension and improve the solubility of hydrophobic compounds (Abdoli et al., 2025). Biosurfactant activity plays a role in biofilm formation and also in biofilm detachment (Ward, 2010). Biofilms are microbial cells that are irreversibly attached to a substratum or an interface embedded in a matrix of extracellular polymeric substances produced by them (Dheilly et al., 2010).

Production of biosurfactants occurs predominantly on hydrophobic substrates such as petroleum hydrocarbons (Ward, 2010). Recently, 55 biosurfactant producers were identified and characterized in North Atlantic Canada. Those belonged to 8 genera, including *Alcanivorax, Exiguobacterium, Halomonas, Rhodococcus, Bacillus, Acinetobacter, Pseudomonas*, and *Streptomyces* (Cai et al., 2014). Biosurfactant production, mediated by the RhlA enzyme, plays a role in the maintenance of the channels within the biofilm and occurs in microcolonies early in biofilm development (Ward, 2010).

Among biosurfactants, glycolipids and glycopeptides with lower molecular weights are generally more effective in reducing surface and interfacial tensions, whereas high molecular weight compounds such as amphipathic polysaccharides, lipopolysaccharides, and lipoproteins serve as stabilizers for oil-in-water mixtures (Lin, 1996). Microbes access the hydrocarbon either via direct interaction with the particulate or liquid droplet hydrophobic substance and interaction with pseudo-solubilized oil or molecules in the form of a surfactant-generated oil-in-water/water-in-oil emulsions, or the surfactant secreted by microbes promotes emulsion formation and facilitates cellular uptake of the hydrophobic substrate (Ward, 2010).

Generally, both monocultures and mixed microbial cultures can be applied in bioremediation; however, higher hydrocarbon degradation rates are often achieved using enrichment consortia isolated from the contaminated environment targeted for restoration (Juhasz et al., 2000). Nonetheless, these naturally occurring surface-active microbial metabolites provide environmentally friendly alternatives to their synthetic counterparts (Abdoli et al., 2025). Hence, bioremediation using biosurfactant-producing oil-degrading autochthonous bacteria serves as a promising green clean-up alternative (Lee et al., 2018).

This study aims to improve understanding of biosurfactant production by cold-adapted marine microbial consortia, which may play a key role in enhancing hydrocarbon bioavailability and influencing oil fate in Arctic environments. Despite the ecological importance of biosurfactants in cold-oil interactions, their functional performance under polar conditions remains insufficiently characterized, particularly in enrichment cultures representing natural microbial communities.

## 2 Methodology

### 2.1 Field sampling at the Churchill Marine Observatory (CMO)

The Churchill Marine Observatory is a multidisciplinary research facility designed specifically for studying oil spills. It features two mesocosm outdoor pools that allow for controlled and experimental investigations in both freshwater and seawater environments contaminated with hydrocarbons. As part of collaborative research conducted by the Centre for Earth Observation Science (CEOS) at the University of Manitoba aiming at investigating the oil behavior in sea ice and the microbial response to such contamination, a total of 114 liters of Churchill diesel oil was added to the experimental pool on December 3^rd^, 2024, followed by systematic sampling of both the sea ice and seawater over the next two months. Approximately 55 days after the oil contamination, light brown aggregates were observed on the surface of the experimental pool, which are referred to as marine oil snow (MOS) (Figure 1).

**Figure 1.**
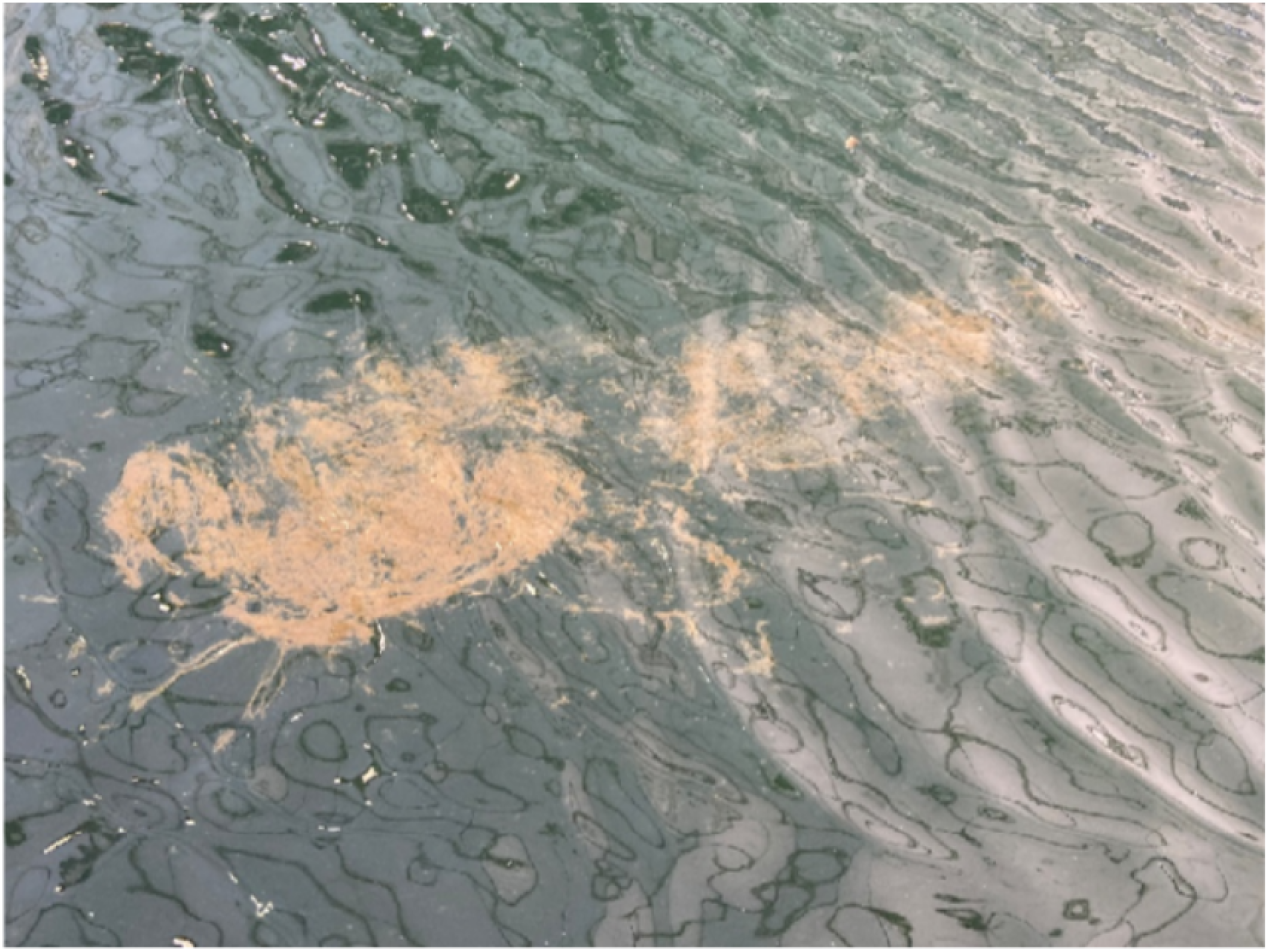
MOS aggregates observed on January 27, 2025, at the surface of the experimental pool in the Ocean–Sea Ice Mesocosm (OSIM) facility at the Churchill Marine Observatory, ∼55 days after the addition of Churchill diesel fuel.

### 2.2 Bacterial cultivation

The MOS aggregates were collected in aseptic amber glass jars and cultured on Bushnell–Haas (BH) agar (plus 3% w/v NaCl) with the motor oil spotted on a sterile filter paper and placed on the lid to use as the sole carbon source for bacterial growth (Malavenda et al., 2015). Cultures were maintained at 4 °C for a month in the dark. Bacterial enrichment cultures (R, T, A, S25, Q1, Q2, Q3, and Q4) were obtained by the streak plate method.

### 2.3 Bacterial growth assessment

Bacterial cultures that could grow with motor oil as the added hydrocarbon substrate were transferred to BH mineral medium (plus 3% w/v NaCl) and allowed to grow aerobically for a month at 4 °C. The composition of the BH medium (g L^−1^) used in this study was: MgSO_4_ (0.20), CaCl_2_ (0.02), KH_2_PO_4_ (1.00), K_2_HPO_4_ (1.00), NH_4_NO_3_ (1.00), and FeCl_3_ (0.05). The pH was adjusted to 7.0, and the medium was autoclaved. Bacterial growth was compared against the respective negative controls, which had no added hydrocarbon substrate.

### 2.4 Screening of biosurfactant producers

Bacterial cultures that grew in the presence of hydrocarbons were screened for their ability to produce surface-active compounds such as biosurfactants (BS) and bioemulsifiers (BE). From each culture, an aliquot of 100 µL was transferred to an Erlenmeyer flask containing 100 mL of BH mineral medium (plus 3% w/v NaCl) and 2% (v/v) motor oil and grown under shaking at 110 rpm for 1 week at 20 °C and 4 °C (Malavenda et al., 2015).

#### 2.4.1 Emulsification index test (E_24_)

The ability of the bacterial cultures to form stable emulsions was assessed using the E_24_ test. After 1 week of growing bacterial consortia under shaking, the medium was centrifuged at 4200 rpm for 40 minutes to obtain the cell-free supernatant. A small volume (2 mL) of each supernatant was added to a flat-bottom glass vial that contained an equal volume of kerosene as the hydrocarbon source. The solution was vigorously vortexed for 1 minute and allowed to stand for 24 hours. The total height of the mixture and the height of the emulsion layer were noted (Malavenda et al., 2015). The following equation was used to calculate the emulsification index of the bacterial isolates, where *h*_*emulsion*_ and *h*_*total*_ are the height of the emulsion layer and the total mixture, respectively. The test was done in triplicate.

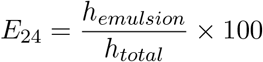

#### 2.4.2 Blue agar plate method

The blue agar plate method is a semi-quantitative assay for the detection of glycolipid biosurfactants or other anionic surfactants such as rhamnolipids (Walter et al., 2013). The blue agar plates were prepared with BH media that were supplemented with peptone 2% (w/v) and cetyltrimethylammonium bromide (CTAB: 0.5 mg/mL)–methylene blue (MB: 0.2 mg/mL) and 3.5% (w/v) NaCl. Inoculum from enrichment cultures was spot inoculated in duplicate onto wells prepared on blue agar plates and incubated at 20 °C for one week. Appearance of dark blue halos around the spotted colonies was considered an indication of surfactant production (Satpute et al., 2008).

#### 2.4.3 Oil displacement method

This method is used to determine the ability of biosurfactants to reduce surface tension and displace oil (Degang et al., 2023). A total of 40 mL of distilled water was added to a Petri dish, followed by the addition of 100 µL of crude oil to form a thin layer on the surface. Then, 10 µL of cell-free supernatant from each liquid enrichment culture was carefully placed onto the oil layer. Formation of a clear zone was considered an indication of biosurfactant production. The diameter of the resulting clear zone was measured to assess biosurfactant activity (Trudgeon et al., 2020). Clear zone diameters were determined by measuring two perpendicular diameters across each zone after 2 minutes and calculating the average. Tween 80 was used as the positive control.

### 2.4.4 Extraction of crude biosurfactants

Putative biosurfactants produced by each bacterial enrichment culture were enriched by acid precipitation. While this simplified procedure enriches for amphiphilic molecules like biosurfactants, it may also co-extract other cell components. A volume of 6 N HCl was added to the cell-free supernatant until pH 2 was achieved, and samples were stored at 4 °C overnight for precipitation. The solution was then centrifuged at 4200 rpm for 1 hour to collect the precipitate, resuspended in distilled water, and neutralized. The crude precipitates were then subjected to a methanol–chloroform extraction (1:1), followed by rotary evaporation at 40 °C. The pellet was collected in pre-combusted (450 °C for 5 h) glass vials, which were dried in a drying oven at 80 °C for 72 h (Phulpoto et al., 2020).

As residual oil did not fully evaporate during oven drying, the dried material was subsequently dissolved in chloroform and filtered onto pre-combusted, pre-weighed GF/F filters. Filters were then dried at 80 °C for an additional 72 h, after which the dry weight of the crude biosurfactant was determined gravimetrically using the following formulae:

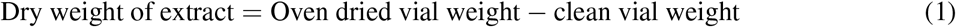

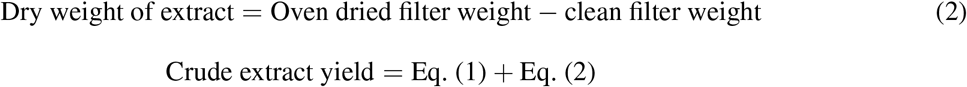

#### 2.4.5 Bacterial phylogenetic characterization

Genomic DNA of the enrichment cultures was extracted using the PowerSoil DNA isolation kit (Qiagen, CA, USA) following the manufacturer’s instructions. The V4–V5 hypervariable regions of the 16S rRNA gene were amplified using the 515F (GTGYCAGCMGCCGCGGTAA) and 806RB (GGACTACNVGGGTWTC-TAAT) primer pair (CD Genomics, New York, USA) and were sequenced using paired-end (2 × 250 bp) Illumina NovaSeq (CD Genomics, New York, USA). Quality filtering of the reads was performed using the DADA2 package in R (Callahan et al., 2016), resulting in 250 k reads with an average depth per sample of 31 k reads. Sequences were aligned using the SILVA v138.2 reference database for taxonomic classification (Quast et al., 2013).

## 3 Results

### 3.1 Qualitative assays of the biosurfactants

#### 3.1.1 Foaming

Surfactants can foam, characterized by bubbles visible through the liquid and on the gas-liquid interface (Satpute et al., 2010). All eight cultures exhibited visible foaming for approximately 2 hours after incubation on a shaker at 110 rpm, at both 20 °C and 4 °C (Figure 2). This observation suggests biosurfactant production by the bacterial enrichment cultures.

**Figure 2.**
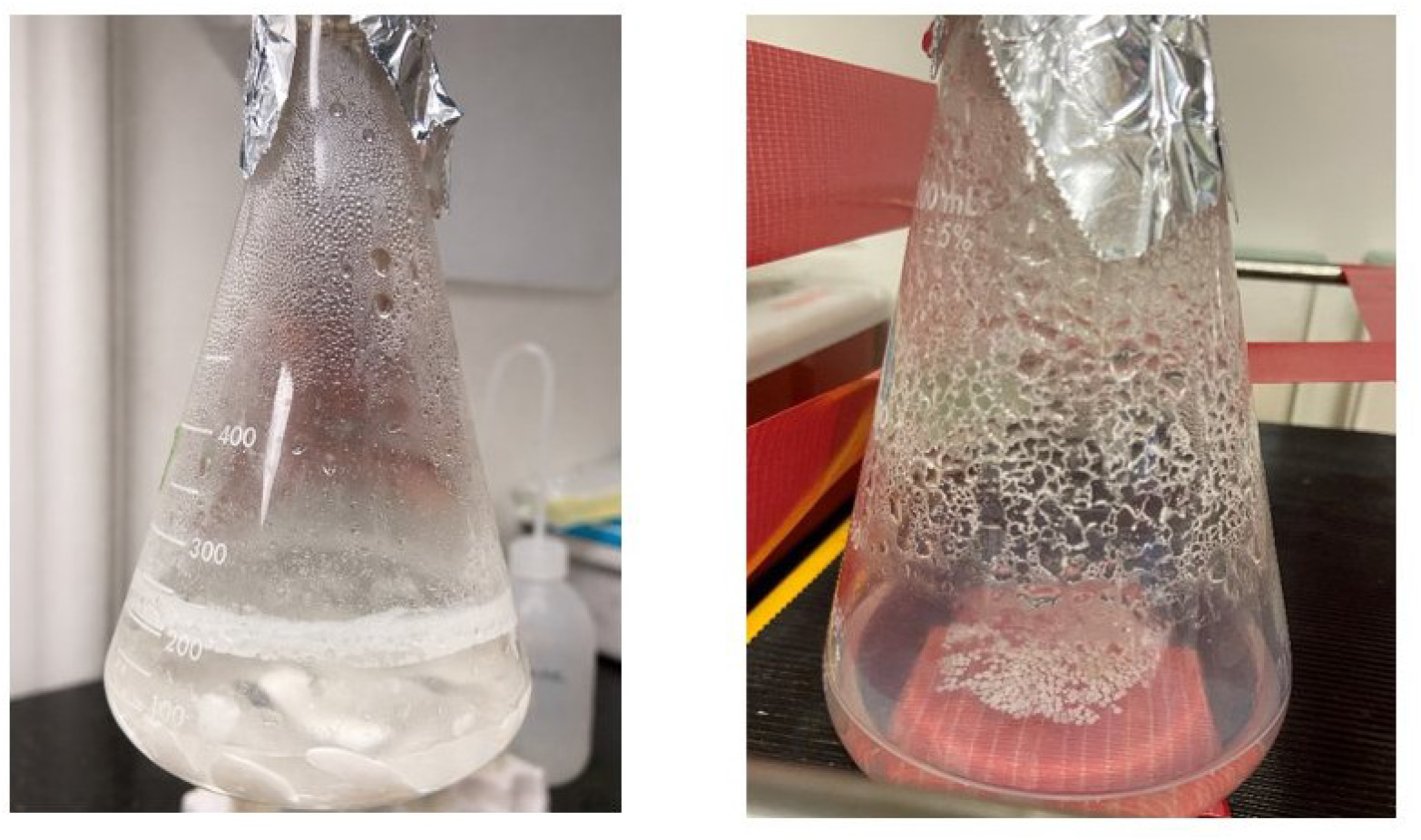
Foaming properties observed in liquid enrichment cultures at 4 °C. The culture consisted of 100 mL of BH mineral media, 2 mL of motor oil, and 100 µL of bacterial inoculum.

#### 3.1.2 E_24_ test

The ability of bacteria to produce stable emulsions was analyzed using the E_24_ test. The emulsification index (E_24_ %) indicates the strength of the biosurfactant. The samples exhibited varying emulsion indices when produced at different temperatures, with all samples tested at a neutral pH of 6–8. All samples, except for A, demonstrated stable emulsions at 4 °C (Figure 3 & Figure S3). At 20 °C, only three samples (R, S25, and Q1) showed stable emulsions that were 16.1%, 3.2%, and 6.5%, respectively. R demonstrated notably higher emulsion stability at 20 °C (16.1%). None of the above bacterial consortia can be considered efficient bioemulsifiers under the above environmental conditions since the E_24_ values were relatively low.

**Figure 3.**
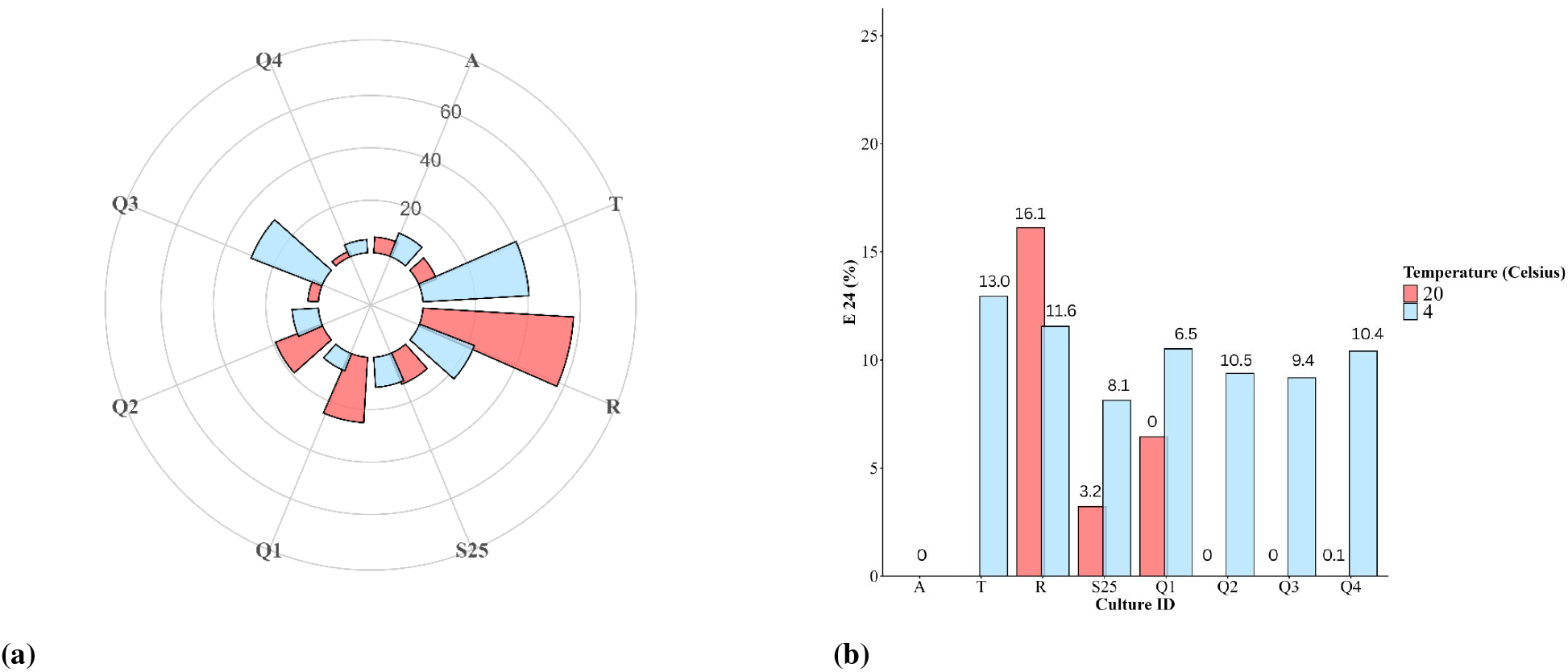
Oil displacement assay of the biosurfactants at 20 °C (red) and 4 °C (blue) (a); E_24_ % of the biosurfactants at 20 °C (red) and 4 °C (blue) (b).

#### 3.1.3 Oil displacement method

R demonstrated the highest biosurfactant activity (57.5 mm) by pushing the oil layer to the edges, while T, Q1, and Q3 exhibited moderate biosurfactant activities. However, the activity varied at different temperatures (Figures 3 & 4).

**Figure 4.**
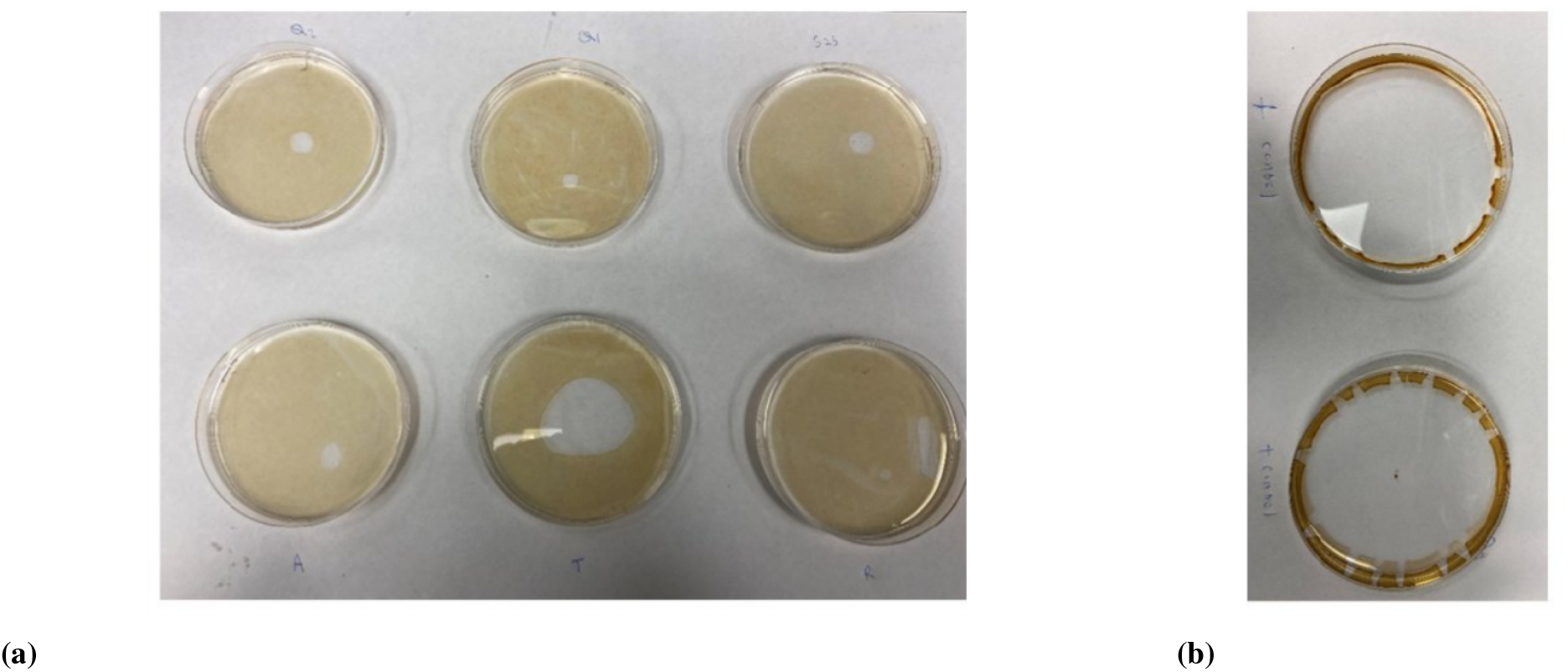
Oil displacement assay of biosurfactants tested on distilled water with crude oil on the top at 4 °C. The samples are arranged from left to right as follows: Q2, Q1, S25 (top row), A, T, and R (bottom row) (a), while Tween 80 serves as the positive control (b).

#### 3.1.4 Blue agar test

Enrichment cultures A, R, and S25 showed positive results in the blue agar assay, indicating potential rhamnolipid production (Figure 5).

**Figure 5.**
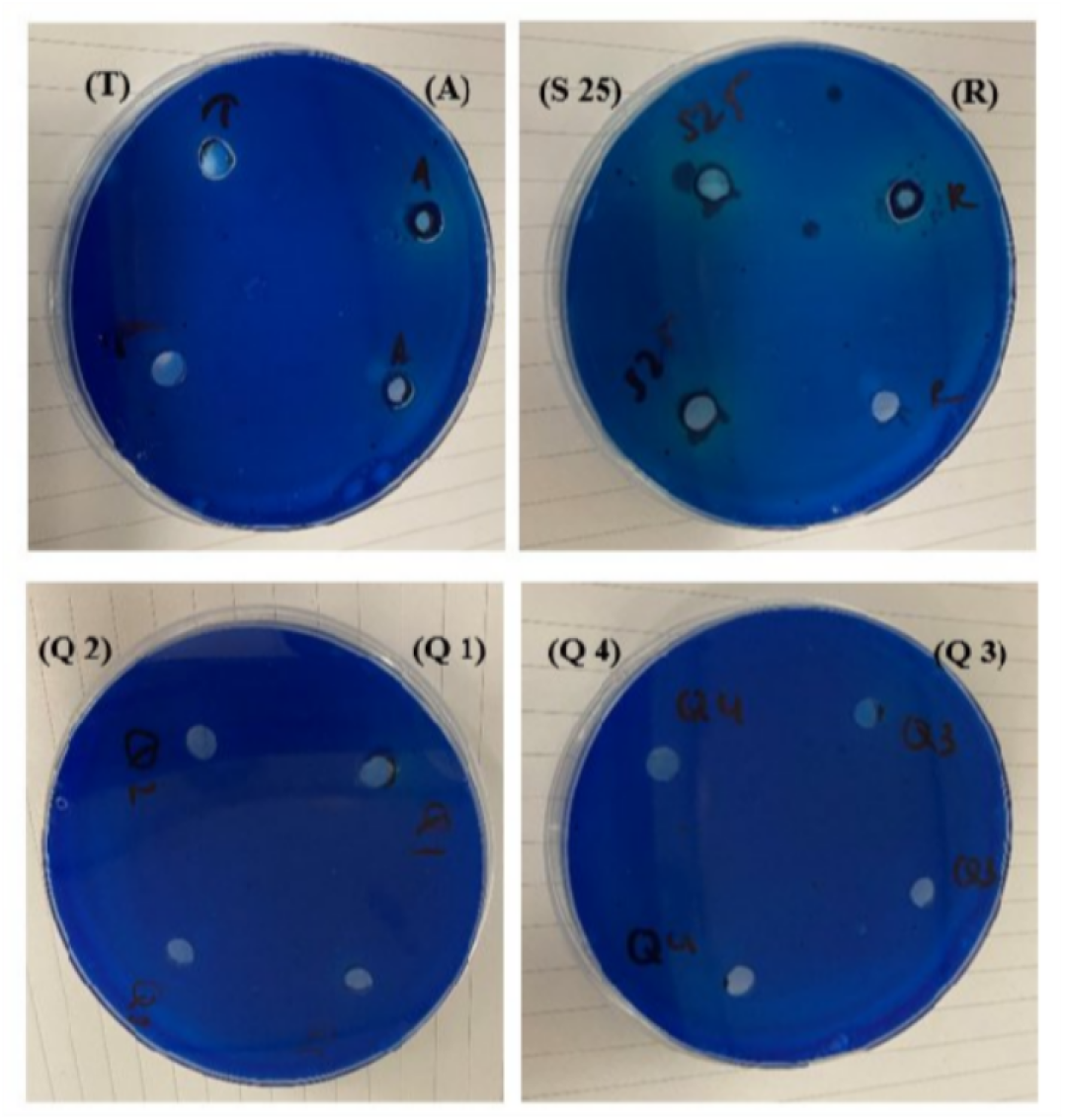
Blue agar test for biosurfactants tested at room temperature (20 °C), showing blue halo zones around A, S25, and R inoculated spots.

#### 3.1.5 Acid extraction yield

A Mann–Whitney U test revealed a significant difference between yields of crude extractions obtained at 4 °C and 20 °C (*W* = 56, *p* = 0.014), indicating a consistent directional trend with decreasing temperature. The magnitude of yield changes varied considerably among samples, with some exhibiting substantial increases at 20 °C (Q1) (Figure 6 & Figure S1). These patterns suggest strain-specific and potentially cold-adaptive production of putative biosurfactants.

**Figure 6.**
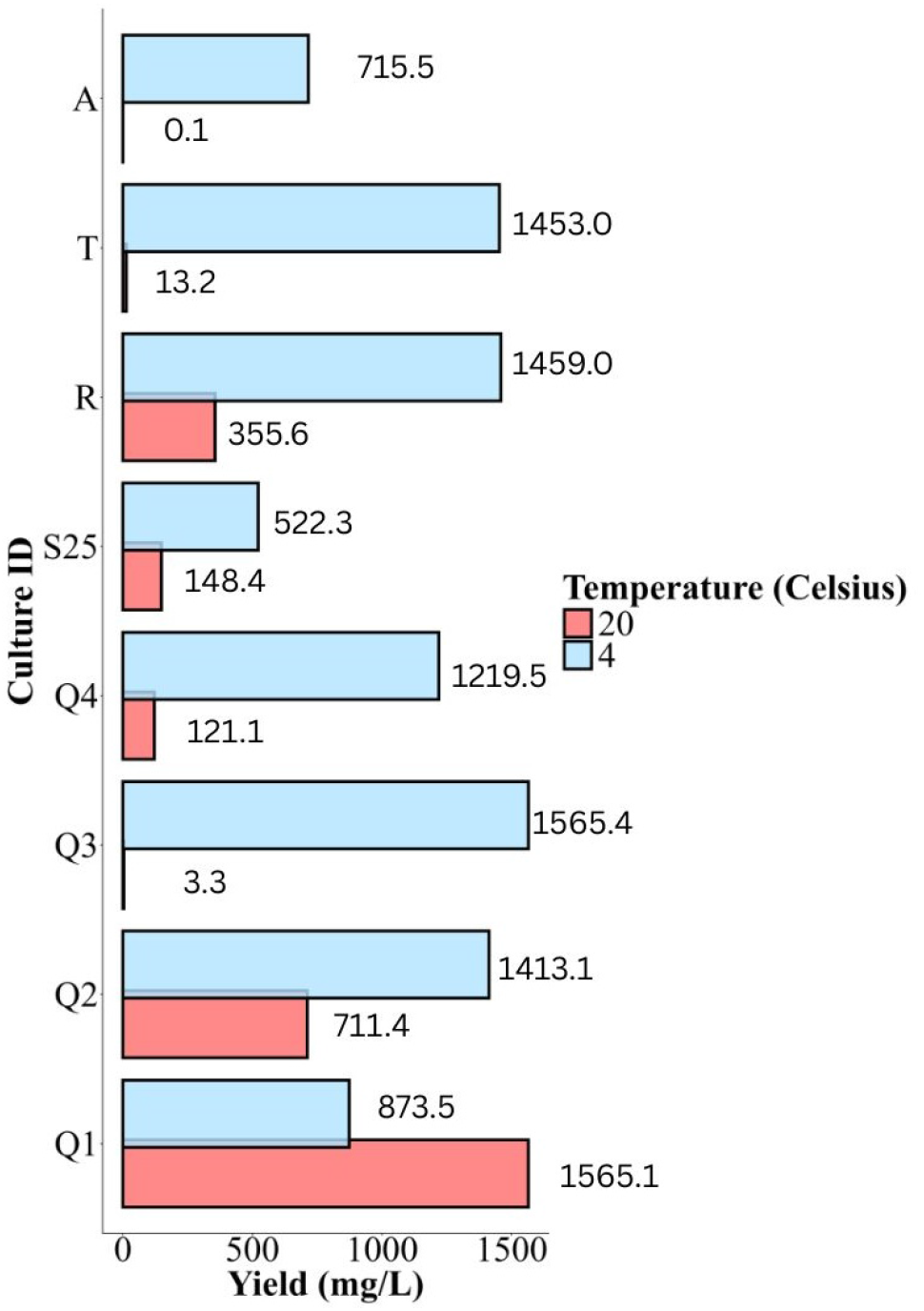
Acid extract yield from bacterial enrichment cultures at two temperatures, 20 °C (red bars), 4 °C (blue bars).

#### 3.1.6 Phylogeny of surfactant producers

Across all enrichments (A, Q1–Q4, R, T, and S25), *Pseudoalteromonas* accounted for the majority of the total relative abundance, ranging from 73% to 98% (Figure 7) with the highest relative abundances observed in cultures S25 (98%) and T (93%), indicating consistent enrichment of this genus under cold, motor oil-based growth conditions.

**Figure 7.**
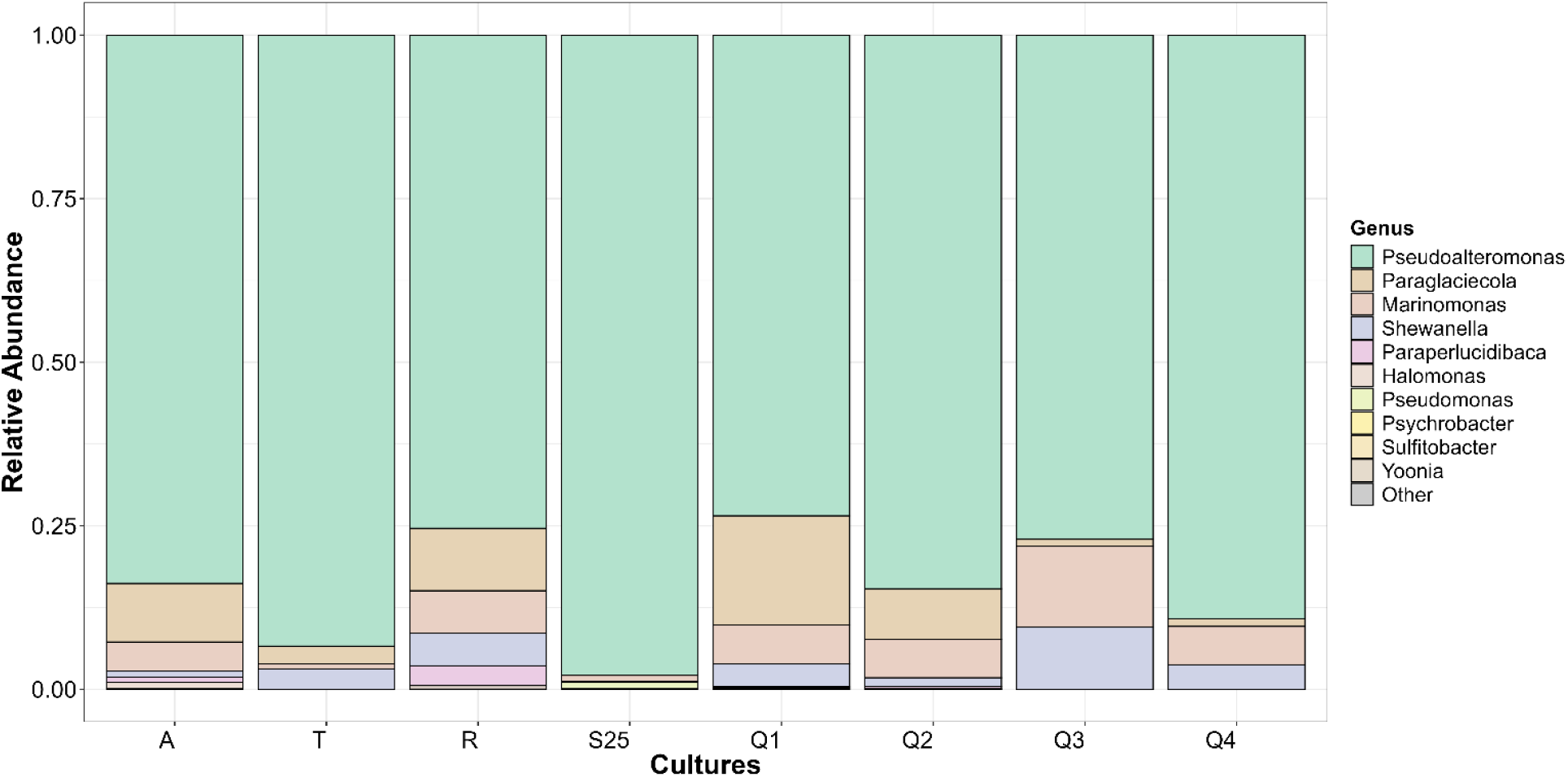
Bacterial community composition of enrichment cultures (A, Q1–Q4, R, T, and S25) incubated at 4 °C with motor oil as the sole source of carbon.

In addition to *Pseudoalteromonas*, several other known surfactant-producing marine genera were detected at lower relative abundances. *Paraglaciecola* was the most prominent secondary contributor, particularly in Q1 (17%), R (10%), and Q2 (8%). *Marinomonas* was also present in multiple enrichments, with its highest relative abundance observed in Q3 (12%). Other genera, including *Shewanella, Paraperlucidibaca, Halomonas, Psychrobacter*, and *Sulfitobacter*, were detected sporadically and generally comprised less than 5% of the community in each enrichment culture.

Community composition varied among enrichments in terms of relative contributions of secondary taxa; however, overall community diversity was low, particularly in cultures T and S25, which exhibited near-exclusive dominance by *Pseudoalteromonas*. These patterns indicate strong and consistent selection for a limited subset of surfactant-producing bacteria under cold, exogenous carbon enrichment conditions.

## 4 Discussion

A total of eight bacterial enrichment cultures were screened for the ability to produce biosurfactants on Bushnell–Haas media in the presence of motor oil as the hydrocarbon source. Across all enrichment treatments (A, Q1–Q4, R, T, and S25), *Pseudoalteromonas* dominated the microbial community, representing 73–98% of the total relative abundance. The highest proportions observed in cultures S25 (98%) and T (93%) indicate strong and consistent selection of this genus under cold, motor oil–amended conditions. This pattern suggests that the enrichment strategy imposed significant selective pressure favoring taxa capable of surviving and proliferating under hydrocarbon stress at low temperatures. Members of this genus are members of the ‘generalist’ hydrocarbon-degrading community and are known for their metabolic versatility (Gutierrez et al., 2013), surface-associated lifestyles (Thomas et al., 2008), and potential production of extracellular compounds (Suja et al., 2017) that may facilitate hydrocarbon interaction and emulsification. The dominance of *Pseudoalteromonas* therefore indicates its potential ecological role in biosurfactant-associated processes and hydrocarbon adaptation under cold conditions.

Although biosurfactants produced at 4 °C demonstrated measurable emulsification activity, none reached the E_24_ ≥ 40% benchmark commonly used to classify strong bioemulsifiers. These results indicate that cold-temperature conditions supported biosurfactant production to some extent, but the emulsifying performance remained moderate.

The methylene blue (MB), cetyltrimethylammonium bromide (CTAB) agar plate assay (Pinzon and Ju, 2009) was used for rapid screening of rhamnolipid-producing cultures. Positive halo formation was observed for enrichment cultures A, R, and S25, indicating production of anionic surfactant(s), possibly including rhamnolipids.

The oil displacement assay performed on enrichment cultures revealed clear temperature-dependent dif-ferences in biosurfactant activity. The highest oil displacement was observed for R produced at 20 °C (57.5 mm), followed by T produced at 4 °C (40.5 mm). Although T was overwhelmingly dominated by *Pseudoalteromonas* (93%), enrichment culture R exhibited a more diverse community structure, comprising *Pseudoalteromonas* (75%), *Paraglaciecola* (10%), *Marinomonas* (7%), and *Shewanella* (5%). The superior oil displacement observed in R, produced at 20 °C, may reflect enhanced microbial metabolic activity at this temperature, which can promote synergistic interactions such as complementary metabolic pathways, cooperative EPS production, and cross-feeding mechanisms that improve hydrocarbon solubilization.

Although all enrichment cultures exhibited broadly similar microbial community compositions, their relative abundances differed, and their functional responses to biosurfactant screening assays varied considerably. Some cultures showed higher oil displacement and greater biosurfactant yield, whereas others did not, despite taxonomic similarity. This suggests that functional performance is not determined solely by bacterial community, but rather by the abundance of key taxa that influence the potential synergistic interactions within the consortium.

Importantly, biosurfactant production can vary even among strains within the same species, meaning that genetically similar isolates may still exhibit markedly different levels of biosurfactant activity (Suthar and Nerurkar, 2016; Aboelkhair et al., 2022). Therefore, even minor shifts in strain-level composition undetectable at higher taxonomic resolution may explain the observed differences in oil displacement and yield among our enrichment cultures. These findings highlight the importance of considering intraspecific diversity and functional heterogeneity when evaluating biosurfactant-mediated oil degradation processes.

Although both T and R exhibited higher biosurfactant yields at 4 °C, their oil displacement activity did not consistently correspond to yield values. For example, isolate R produced a higher acid extract yield at 4 °C (1459 mg L^*−*1^) compared to 20 °C (355.6 mg L^*−*1^); however, oil displacement was greater at 20 °C (57.5 mm) than at 4 °C (22.5 mm). Similar temperature-dependent discrepancies between acid extract yield and oil displacement activity were observed in enrichment cultures Q4 and Q2. These results indicate that acid extract yield alone does not necessarily predict their performance, suggesting that differences in chemical composition, structural properties, or community interactions may influence functional effectiveness. Thus, temperature may affect not only production levels but also the physicochemical characteristics and emulsification efficiency of the biosurfactants produced, as demonstrated in previous studies (Ortiz et al., 2021).

## 5 Conclusion

This study evaluated the biosurfactant-producing potential of eight bacterial enrichments derived from marine oil snow collected at the Ocean–Sea Ice Mesocosm (OSIM) facility at the Churchill Marine Observatory. Biosurfactant production varied significantly with incubation temperature, with enhanced activity observed at lower temperatures. Given that these bacterial enrichments originated from polar environments, their metabolic activity is potentially adapted to low-temperature conditions. Among the enrichments, R exhibited the highest emulsification and oil displacement activity at 20 °C, along with a comparatively high acid extract at 4 °C. Enrichment R also tested positive in the blue agar assay, suggesting the possible production of a rhamnolipid-type biosurfactant. Collectively, these findings highlight the potential of cold-adapted marine bacterial communities associated with marine oil snow to produce biosurfactants under low-temperature conditions. With further characterization and optimization, biosurfactants derived from such strains may represent promising candidates for future bioremediation applications in cold marine environments.

## Data availability

All raw 16S rRNA gene sequence reads generated from this experiment have been deposited at NCBI’s Sequence Read Archive (SRA) under the BioProject PRJNA1176269 (accession numbers SAMN54986504 to SAMN54986511).

## Competing interests

The authors have declared no competing interest.

## Author contributions

Thamali Vidanage – Conceptualization, Data Curation, Formal analysis, Investigation, Methodology, Visual-ization, Writing – original draft, Writing – Review & Editing, Project Administration Durell Desmond – Conceptualization, Methodology, Data Curation R. Eric Collins – Conceptualization, Methodology, Writing – Review & Editing, Validation, Supervision, Funding acquisition, Project Administration

## Acknowledgements

We would like to sincerely thank Dr. Marike Palmer (Department of Microbiology, University of Manitoba) for generously providing laboratory space for culture media preparation and access to chemicals required for this project, and Dr. Feiyue Wang and Debbie Armstrong for generously providing the chemicals used in culture media preparation. We are grateful to Marcos Lemes for providing access to the necessary laboratory instruments for culture media preparation.

We are also thankful to Jocelyn Plouffe for her assistance with purchasing the required chemicals and support with sample preparation. We would like to thank Lisa Oswald and Shiva Lashkari for their assistance with crude biosurfactant extraction. Additionally, we thank Cyril Fredlund for his support during the fieldwork at the CMO.

## Funding

This research work was funded by a Genome Canada LSARP grant to R.E. Collins (GENICE II).

## Supplementary material

### Data tables

**Table S1.**
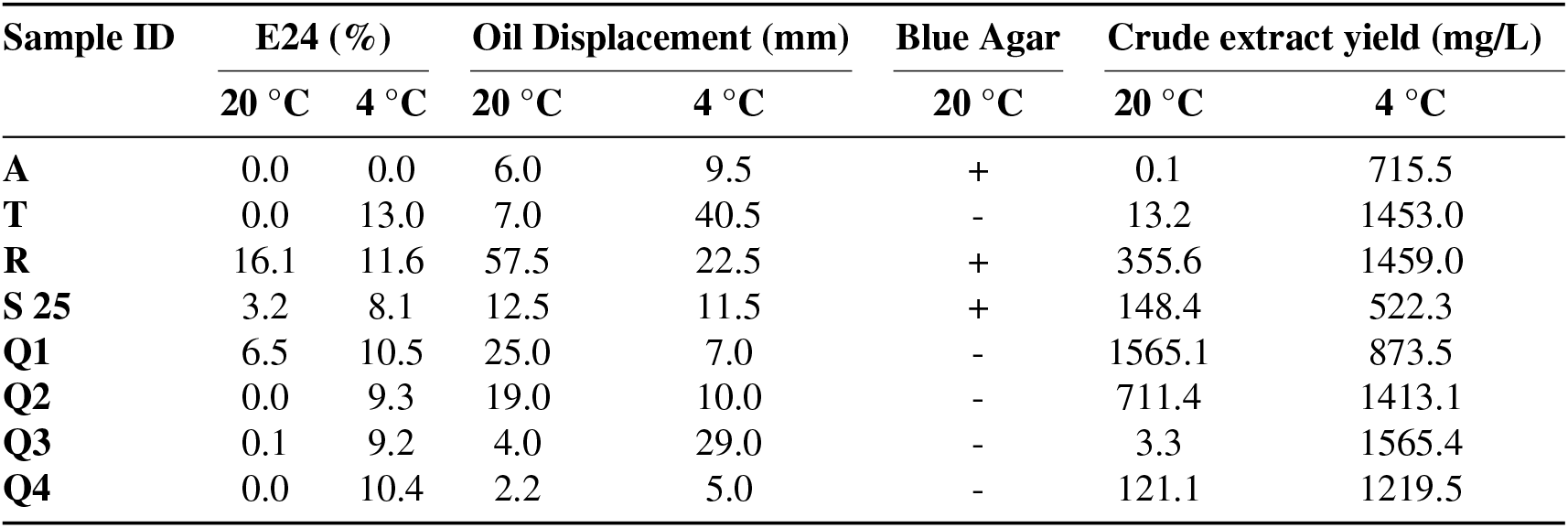
Summary of the biosurfactant screening results for samples (A–Q4) at 20 °C and 4 °C.

### Additional figures

**Figure S1.**
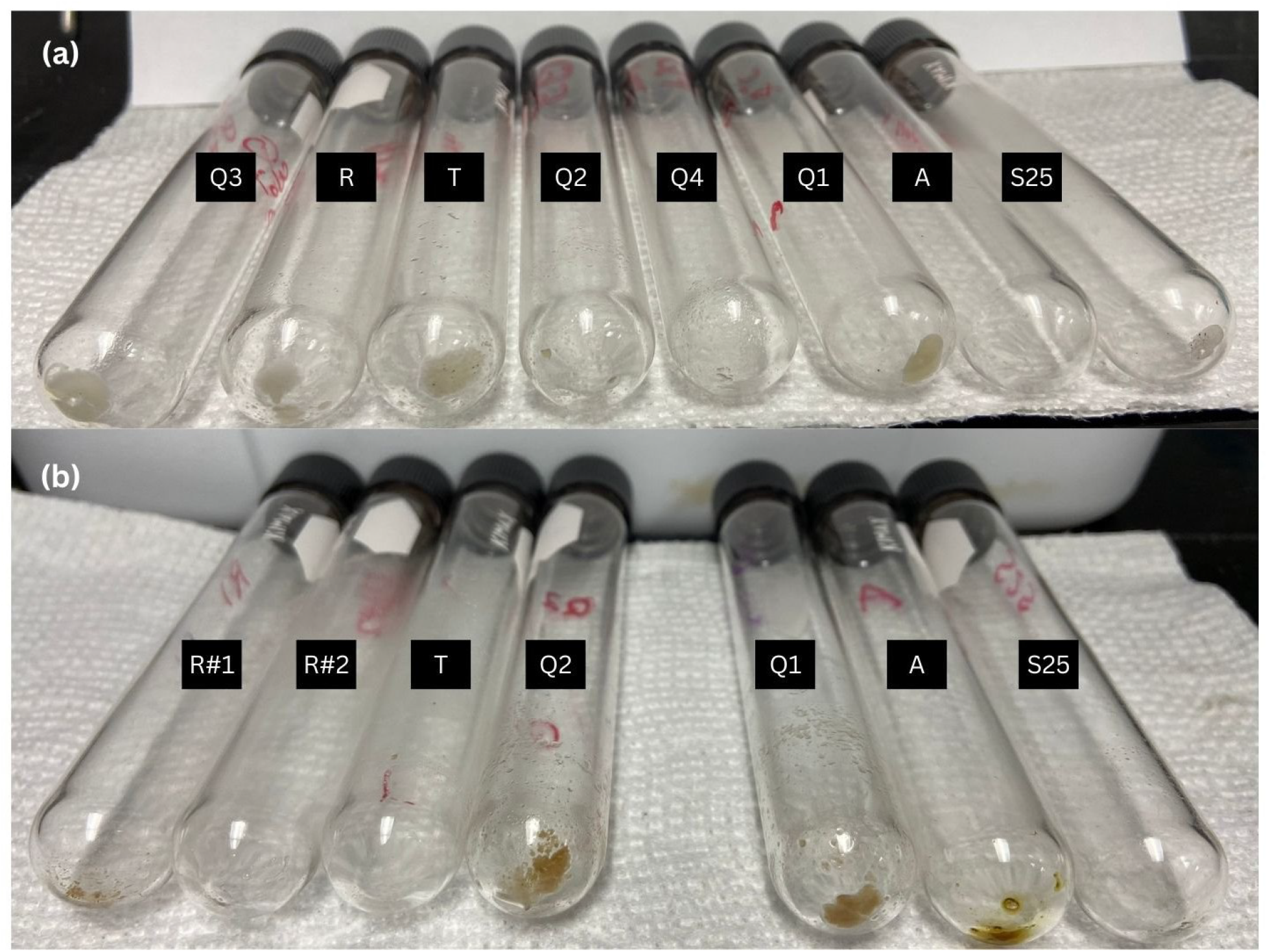
Extracted biosurfactant products obtained from different enrichment cultures incubated at (a) 4 °C and (b) 20 °C. Cultures are arranged from highest to lowest biosurfactant yield based on production at 4 °C. Missing vials in panel (b) indicate samples that were entirely filtered onto pre-combusted GF/F filters for dry weight determination.

**Figure S2.**
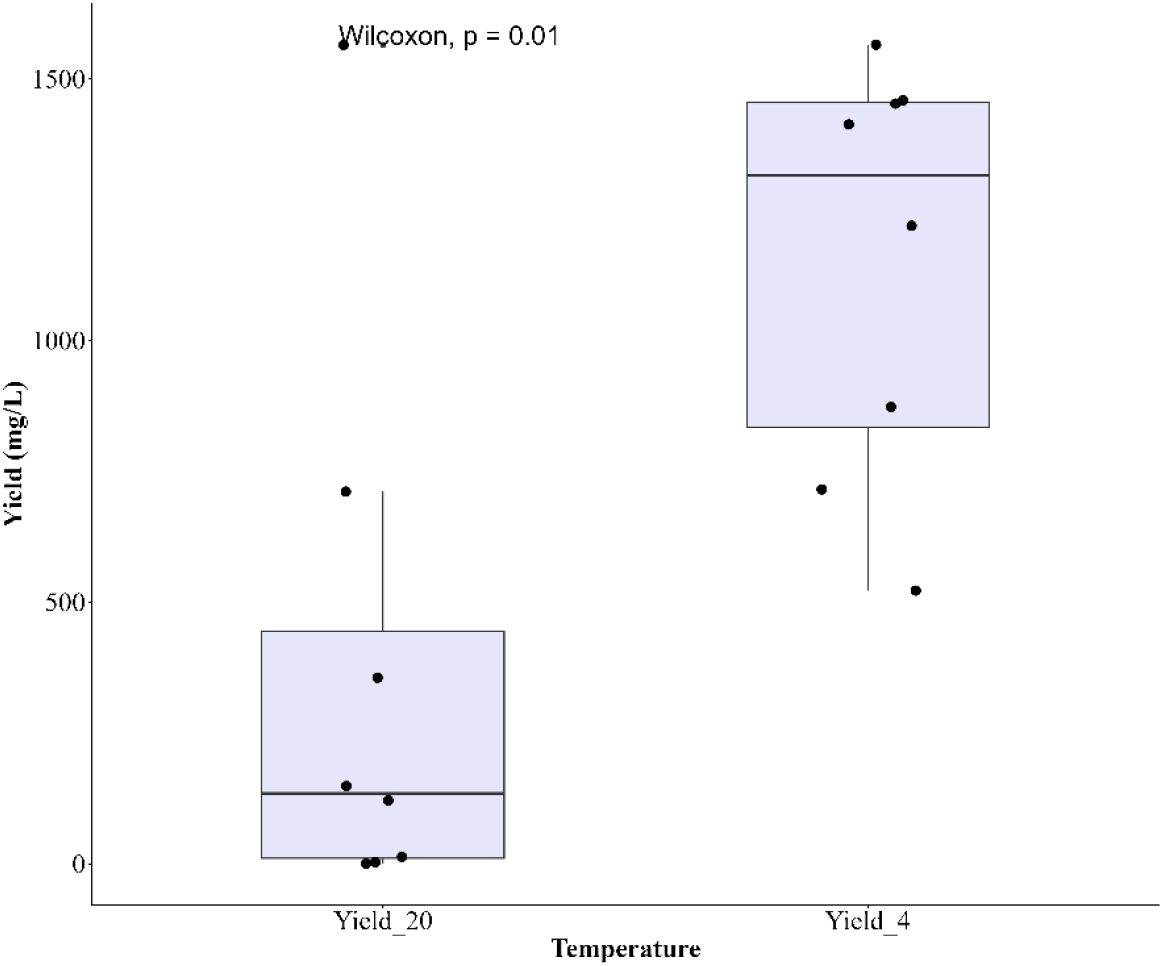
Biosurfactant yield produced at 4 °C and 20 °C by enrichment cultures. Boxes indicate the interquartile range and median, while points represent individual yield measurements. Yield differed significantly between temperatures based on a Mann–Whitney U test (*p* = 0.014).

**Figure S3.**
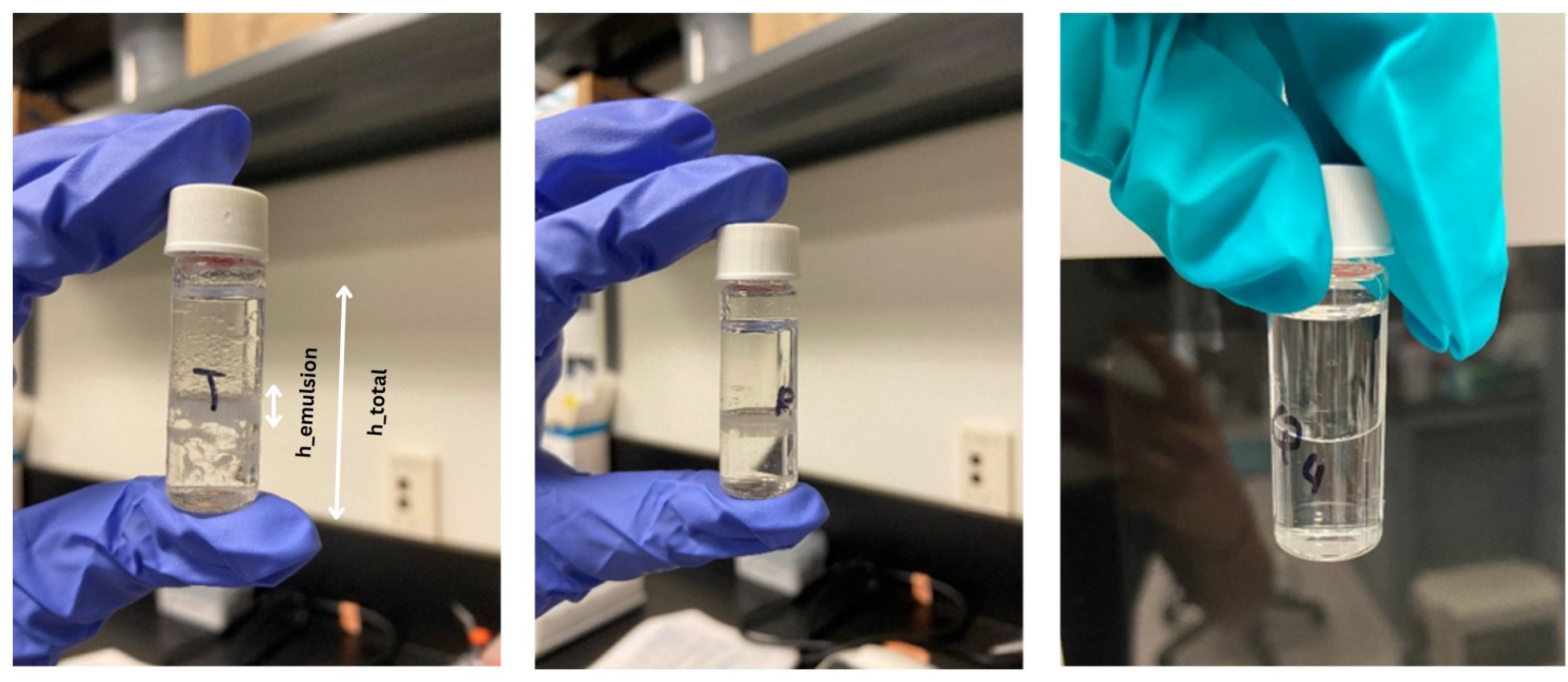
Emulsification index (E_24_) test of cell-free supernatants from enrichment cultures T and R grown at 4 °C, and Q4 grown at 20 °C, demonstrating their ability to form stable emulsions. Visible emulsification was observed in cultures T and R, whereas Q4 showed no detectable emulsification after 24 h.

## Notes

https://www.ncbi.nlm.nih.gov/bioproject/PRJNA1176269

